# A systems biology approach to elucidate the post-translational regulome of coronary artery disease

**DOI:** 10.1101/2022.02.09.479759

**Authors:** Ankit Sharma, Madankumar Ghatge, Vrushali Deshpande, Rajani Kanth Vangala

## Abstract

Coronary Artery Disease is a major killer in India and world at large but the molecular regulators which modulate clinically relevant pathways are not completely understood. This study was aimed at identifying essential post-translational modifications (PTM) regulome network and its master regulator modulating the CAD associated pathways. 995 CAD associated genes were taken from InCardiome database (www.tri-incardiome.org) were analyzed for all possible PTMs. Two important interdependent molecular processes which define the function of a protein are protein-protein interactions and PTMs of which PTMs play regulatory role. Using PTMCode2 we evaluated the co-evolving amino acid residues for important PTMs and found that serine-serine phosphorylation is highly represented combinatorial regulator in these set of proteins. Furthermore, in the CAD associated pathways we again found that serine phosphorylation was dominant player in all the processes of atherosclerosis. In order to identify the master regulator kinase, we further assessed the kinome network associated with CAD and identified 5 most important kinases namely GSK3B, PRKCA, PRKCD, SRC and PRKACA which might modulate clinically important pathways. GSK3B with the highest network parameters (node degree and betweenness centrality) was identified as master regulator and 1 U/l increase of phsophoGSK3B (on a log scale) increased the odds ratio (OR) by 4.07 fold (AUC 0.620) and 6.27 fold (AUC 0.752) upon addition of conventional risk factors (CRFs).

## Introduction

While the impact of the coronary artery disease (CAD) is on rise in India, most of the studies are at infancy with regard to understanding the multi-factorial disease. The pre mature onset of the disease among the Indians translates to highest loss of potentially productive years worldwide [1]. Current treatments for atherosclerosis are based on the various classical risk factors like hypertension, diabetes, hyperlipidemia, smoking and obesity. Concerted efforts are on to reduce the prevalence of these modifiable risk factors. However, many CAD patients do not have any of these identifiable traditional risk factors [2]. Emerging biomarkers (genetic, circulating and sub clinical and clinical) have been shown to improve risk stratification beyond the baseline Framingham model and reclassify individuals into their appropriate risk groups [3]. Studies have shown that post translational modifications (PTMs) are important regulators of protein functions thus also the pathways. Almost 200 thousand PTMs have been identified by studying individual proteins using radiolabeling, western analysis, and mutagenesis [4-6]. Post translational modification of fibrinogen by reactive nitrogen species is important to maintain the pro-thrombotic state and can also be used as risk predictor [7].In case of Pro-Brain Natriuretic Peptide, glycosylation has been shown as a major factor for prediction heart failure studies [8].Therefore, the present study addresses the most important aspect of understanding the disease biomarker’s regulation using integrative methodology of bioinformatics and proteomics. This may further lead to identification of new potential mechanisms and use of PTM level as a risk assessment tool, suggesting the holistic measurement of modulation of pathway regulators as tools in diagnostics and risk prediction of CAD.

Our methodology of identification of important PTMs and its regulators lies in using system biology approach. Information for CAD Genes was taken from literatures, existing data sources and reliable data mining tools using different search terms. PTMs information was also collected for identified CAD genes which were further curated in house. Bioinformatics approach has been implemented to identify top PTMs and associated regulators. Here we have shown the association of serine phosphorylation and its important regulator GSK3b in CAD. Further we validated our findings in patient population from Indian Atherosclerosis Research Study (IARS) cohort [9].

## Methodology

Methodology for this work is shown in figure 1.

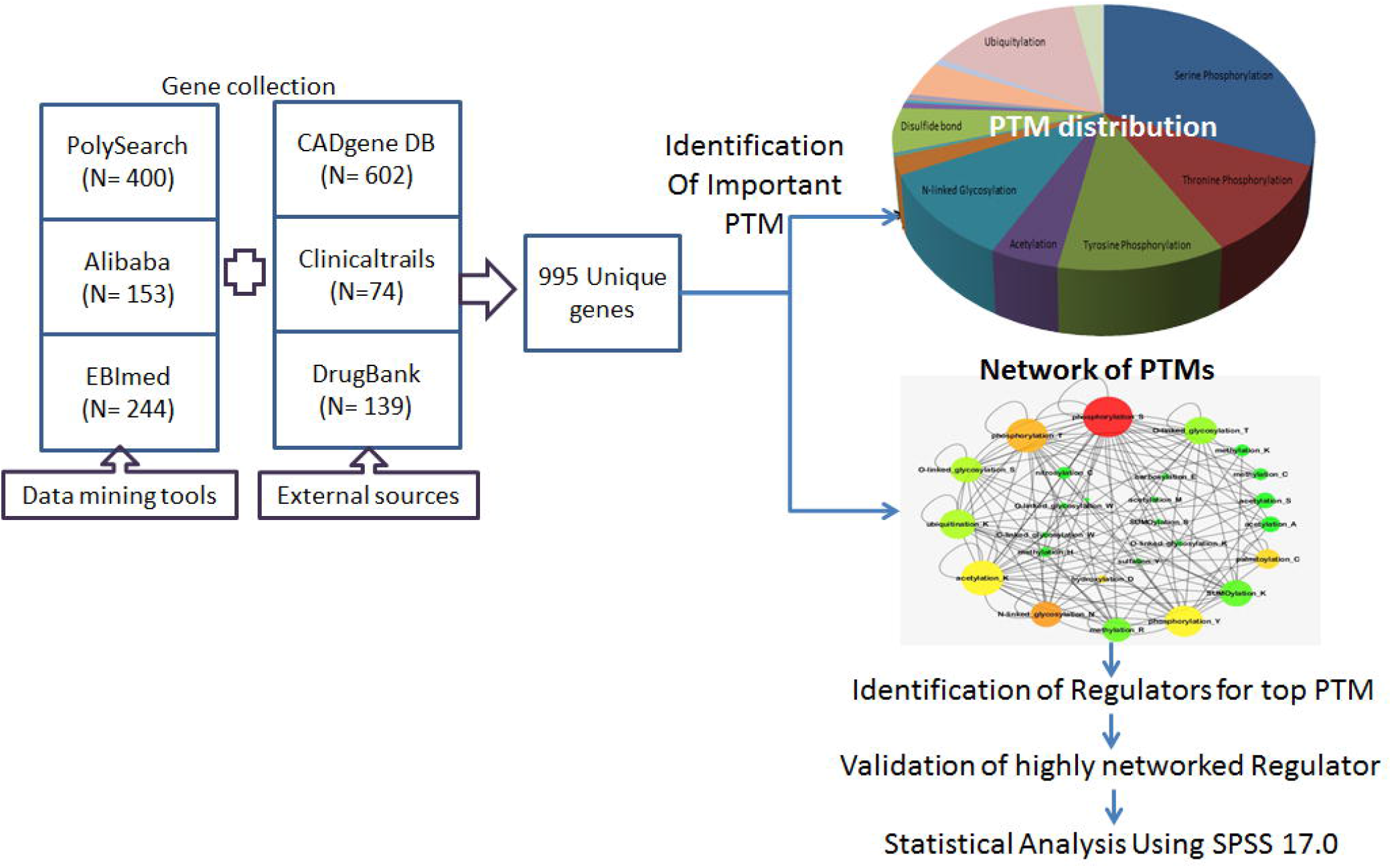

### Identification of CAD associated genes

We selected CAD associated genes from InCardiome database (www.tri-cardiome.com), the information in this database was added by three text mining tools namely PolySearch [10], Ali-baba [11] and EBImed [12] using a search term “coronary artery disease”. The same term was used to extract genes from Clinical trails.gov [13] and DrugBank [14]; also incorporated genes from CADgene database [15] (figure 1). After accumulation of gene list from each tool, all were merged and a unique list of genes was generated by removing redundancy.

### Extraction of PTM and its Master regulators

After getting the unique list of genes, we obtained PTM information from dbPTM [16] and Uniprot [17] individually. Further Perl script was used to get frequency of each PTM to identify highly enriched post translational modification.

### Pairs of PTMs

An increasing amount of research has provided enormous data on the diversity and abundance of reported PTMs therefore it is important to distinguish individual PTMs to be functionally associated. Studies have shown the coevolution of the modified residues of PTMs and also that different PTMs work together for any specific function and protein localization in proteome [18]. We further identify functionally associated PTMs which regulate important pathways in coronary artery disease. We downloaded functionally co-evaluated PTM data from PTMcode2 for our interest of genes [19].

### Network analysis

After collecting data on functionally associated PTMs, we filtered them based on their frequency presented in database and removed redundancy of same pair. This unique data of associated PTMs is used for constructing the network to get topmost PTMs, which works together with other PTMs and regulates the diseased pathways. We used Cytoscape 3.0.1 for the network analysis [20]. We used network topological parameters “degree distribution and betweenness centrality” to show the level of association between PTMs.

### Identification of Important regulator for enriched PTM

We identified serine phosphorylation as a major regulator of our set of genes. It has been well established that phosphorylation or dephosphorylation of proteins can alter their expression level in disease condition [21]. Phosphorylation of protein at any amino acid residue gets regulated by specific group of proteins, including Kinases and phosphatase. We collected human protein kinases interactions information from PhosphoPOINT database [22]. In this database information is divided into four categories: Category 1, interacting protein; Category 2, interacting phosphoprotein; Category 3, known substrate; Category 4; interacting phosphoproteins and known substrate. Results from this step were used for constructing input file for network analysis.

Network analysis was performed and hub kinases were selected using topology parameters such as degree distribution and between ness centrality.

### Validation of Phospho-GSK3b as a master regulator

We have identified serine phosphorylation as an important PTM and 5 kinases as regulators for serine phosphorylation. In the network analysis we identified GSK3b as one of the hub kinase and in order to confirm the association between PhosphoGSK3B (S9) and CAD literature survey was done. Further to validate our approach, a total of 297 CAD affected and 336 unaffected subjects were selected from the ongoing Indian Atherosclerosis Research Study (IARS) cohort. These subjects were enrolled during February 2004 to June 2014. A detailed design of the IARS has been previously described. To describe briefly, IARS participants were primarily recruited from Narayana Health (formerly Narayana Hrudayalaya Hospitals), Bangalore and the Asian Heart Institute and Research Center, Mumbai. Enrollment was done based on stringent inclusion and exclusion criteria which included those having disease onset age of 60 years or less for men and 65 years or less for women. CAD was diagnosed by coronary angiography (>70% stenosis in major vessels or >50% stenosis in two or more smaller coronary vessels) with electrocardiogram (ECG) and coronary artery bypass grafting (CABG) or percutaneous coronary intervention. Clinically unaffected subjects had normal ECG readings. All subjects were free from concomitant infection or major illness at the time of enrollment with informed voluntary signed consent obtained. The entire study protocol and projects were approved by Thrombosis Research Institute Institutional Human Ethics Committee abiding by guidelines of Indian Council of Medical Research and following the principles of Declaration of Helsinki 1975.

## Statistical analysis

Statistical analysis was conducted using SPSS, version 17.0 (SPSS, Inc., Chicago, IL, USA). Results for continuous variables were presented as the mean ± standard error mean (SEM) using Student’s independent t□test. χ2 test was used for testing the association of binary variables such as Hypertension, diabetes, gender, alcohol and smoking habit with CAD. Since Classical Risk Factors (CRFs: Age, Gender, Hypertension, diabetes, smoking and higher lipid levels (LDL, TG, TC) are the backbone for CAD, we consider them as covariates and adjusted during model construction. Subset analysis was also performed for 4 different traditional risk factors (hypertension, diabetes, smoking and alcohol) and difference in expression levels were represented as mean ± standard error of mean (SEM). To assess the association of PhosphoGSK3bwith and without CRFs and to estimate odds ratios (ORs) and 95% confidence intervals, logistical regression analysis was performed. P<0.05 was considered to indicate a statistically significant difference.

To assess the accuracy of discrimination using receiver operating characteristics or C statistics analysis was performed by different models of PhosphoGSK3b: Model1, CRFs alone; Model2, PhosphoGSK3b alone and Model3, PhosphoGSK3b and CRFs combined. The statistical significance between the area under the curve (AUCs) was calculated using the DeLong method [23] in R statistical program, version 3.2.1 (https://cran.r-project.org/) and calibration of the prediction models was tested using Hosmer–Lemeshow test.

## Results and Conclusions

### Identification of PTM sites in genes

As presented in table.1, Total 9273 PTM sites were identified for In-Cardiome genes of which approx. 50% of sites are covered by phosphorylation. We found phosphorylation, ubiquitylation acetylation and N glycosylation to be highly existent PTMs (figure 2a).

**Table 1:** PTMs involved in coronary artery disease

| No. | PTMs | No. of PTM sites |
| --- | --- | --- |
| 1. | Serine Phosphorylation | 2899 |
| 2. | Threonine Phosphorylation | 1006 |
| 3. | Tyrosine Phosphorylation | 1002 |
| 4. | Acetylation | 412 |
| 5. | N-linked Glycosylation | 953 |
| 6. | O-linked Glycosylation | 190 |
| 7. | ADP-ribosylation | 3 |
| 8. | Alkylation | 3 |
| 9. | Amidation | 14 |
| 10. | Carboxylation | 20 |
| 11. | Citrullination | 1 |
| 12. | C-linked Glycosylation | 6 |
| 13. | Disulfide bond | 500 |
| 14. | Hydroxylation | 67 |
| 15. | Methylation | 34 |
| 16. | Myristoylation | 7 |
| 17. | Neddylation | 5 |
| 18. | Nitration | 12 |
| 19. | Oxidation | 4 |
| 20. | Palmitoylation | 41 |
| 21. | Prenylation | 2 |
| 22. | Proteolytic Cleavage | 415 |
| 23. | Sumoylation | 83 |
| 24. | Ubiquitylation | 1335 |
| 25. | Others | 259 |
|  | Total | 9273 |

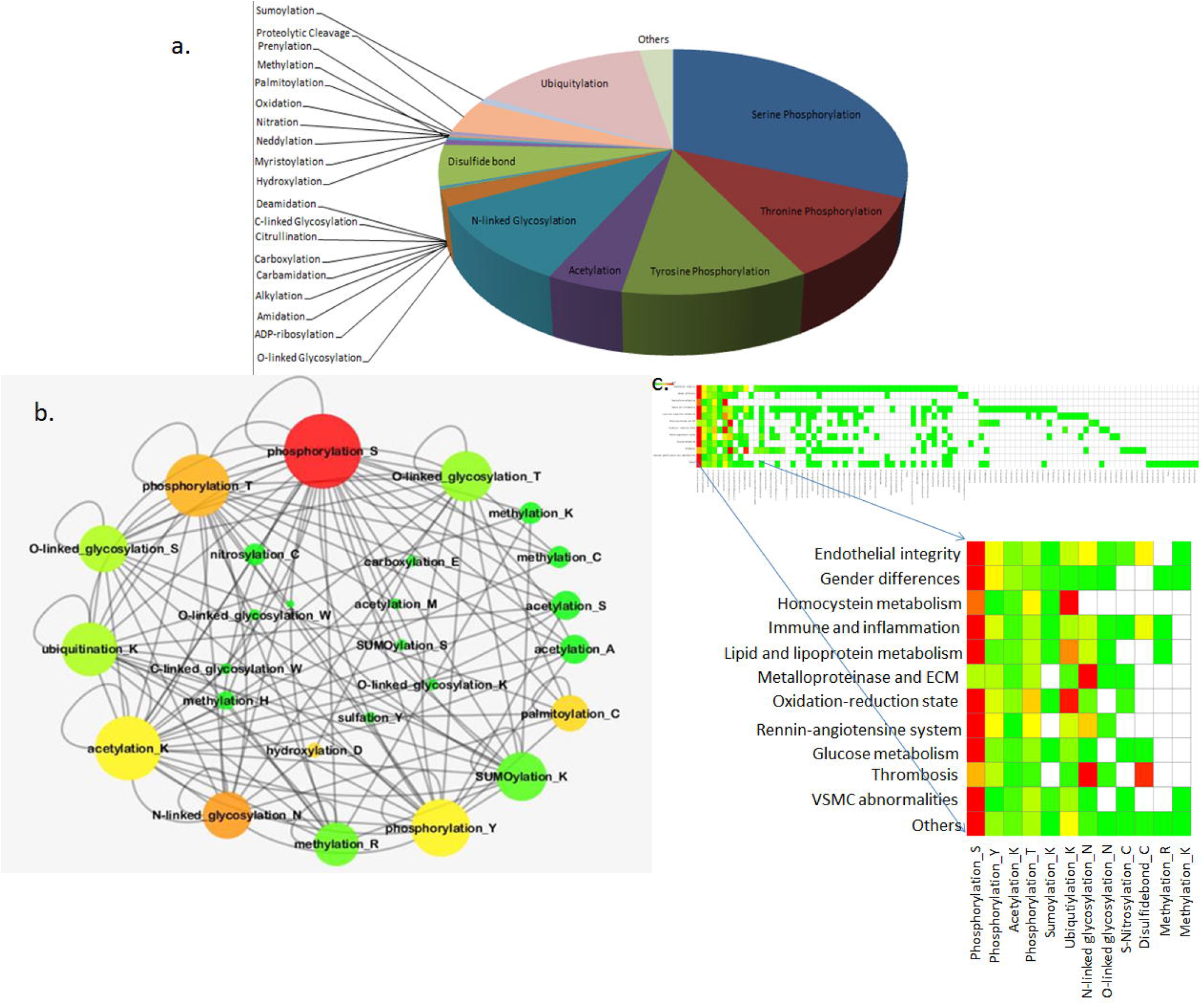

In the second approach of combinatorial regulatory PTMs analysis we identified PTM-PTM combination for each protein (table 2). PTMs prioritization is measured by no. of time two PTMs repeated together in a protein. We identified Phosphoserine - Phosphoserine combination was highly repeated with frequency of 939 and Phosphoserine - Phosphothreonine with frequency of 561 and Phosphoserine - Phosphotyrosin with frequency of 306 at second and third place respectively. After getting the clusters of PTM we constructed a network to identify most repeatedly arising PTM in coronary artery disease. Based on topological parameter -highest degree distribution and between-ness centrality, we selected top four PTMs i.e. Serine phosphorylation, threonine phosphorylation, acetylation and n-glycosylation (figure 2b).

**Table 2:** Frequency of PTM-PTM pairs for CAD proteins.

| Frequency | PTM1 | PTM2 |
| --- | --- | --- |
| 939 | phosphorylation_s | phosphorylation_s |
| 287 | phosphorylation_s | phosphorylation_t |
| 274 | phosphorylation_t | phosphorylation_s |
| 158 | phosphorylation_y | phosphorylation_s |
| 150 | phosphorylation_s | acetylation_k |
| 148 | phosphorylation_s | phosphorylation_y |
| 134 | n-linked_glycosylation_n | n-linked_glycosylation_n |
| 123 | acetylation_k | phosphorylation_s |
| 122 | phosphorylation_y | phosphorylation_y |

In the third approach, we identified the connection between PTM and all the important functional categories for CAD as mentioned in CADgene database. First we fetched all the proteins associated with 12 different functions categories followed by site of PTMs. Frequencies for each PTM were calculated and heatmap was constructed using Matrix2PNG [24] (figure 2c).

By following all three approaches, we found Serine phosphorylation as a highly enriched PTM which might play important role in regulation of genes and associated pathways in coronary artery disease. We identified 286 genes with phosphorylation sites shown in supplementary table 1, which were further used in analysis of identification of important regulators.

### GSK3b as a potential regulator

Out of 286 proteins with serine phosphorylation site of 198 proteins were identified as kinases (figure 3). Using network statistics such as node degree distribution (NDD) and between-ness centrality (BNC), we identified TGFBR1 (NDD:37, BNC:0.180), EGFR(NDD:31, BNC:0.132) GSK3B(NDD:23, BNC:0.08) ABL1(NDD:20, BNC:0.07) and SRC (NDD:16, BNC:0.07) as a hub kinases (supplementary table 1)

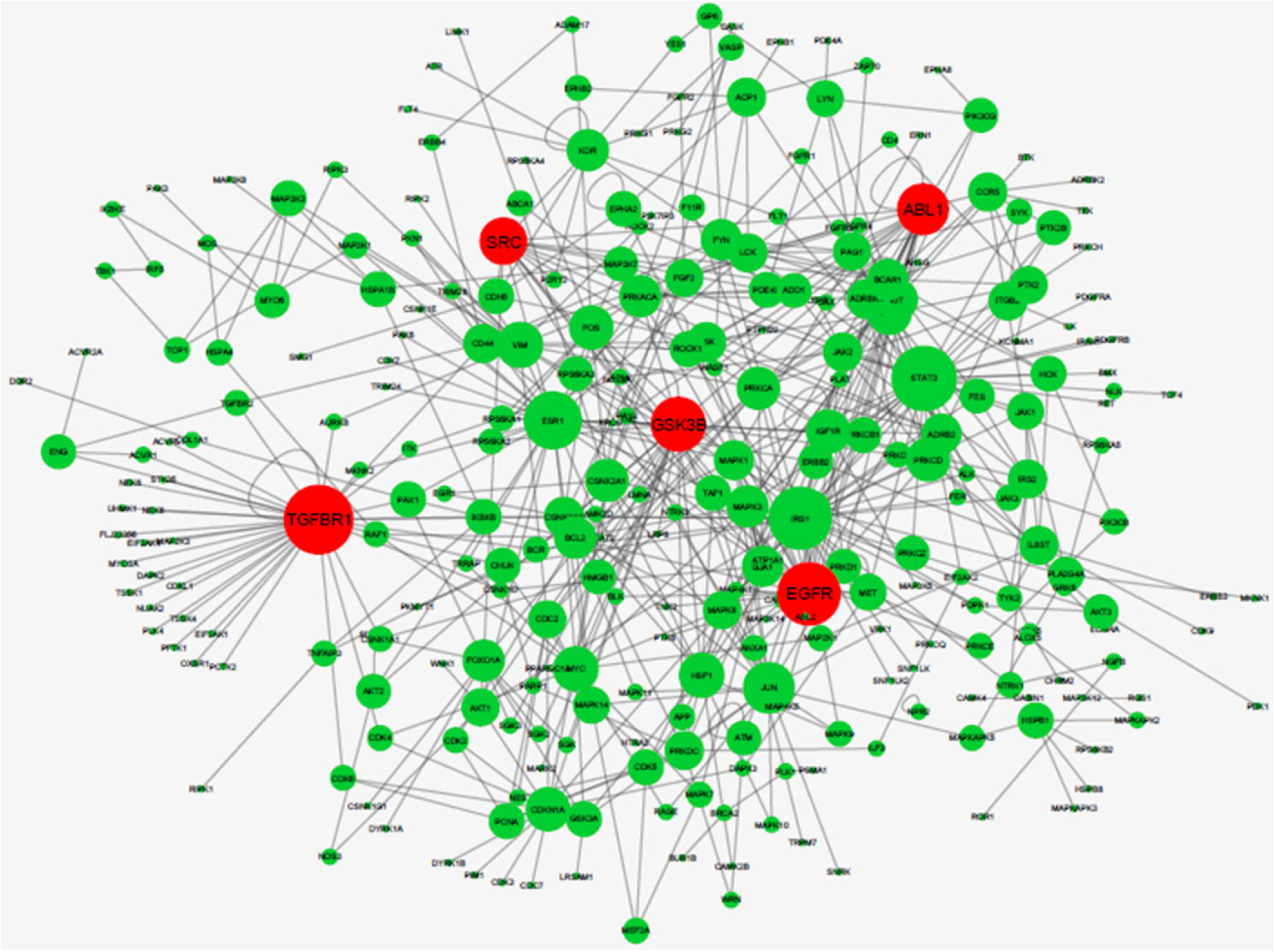

We selected phosphoGSK3B (S9) for further validation in patient sample as it is a downstream protein for all other top kinases (Protein kinase A and Protein kinase B cascade) and very less studied with regards to CAD.In network analysis GSK3Bwas found to be directly connected with genes MYC, JUN, HSF1, CDKN1A, LRP6, HSF1, ESR1 and also had connections with important kinases such as AKT1, PRKACA, PRKCB1, PRKCD, PRKCZ, MARK2, SGK- SGK3, RPS6KA1- RPS6KA3.

### Phospho-GSK3b level and coronary artery disease

Clinical profile of the study population: The baseline characteristics of 632 subjects are shown in table 3. There were 263 males and 72 females in the unaffected and 240 males and 57 females in cad affected group. The mean age in affected group was 42.86 years for males and 47.96 years for females.The Traditional risk factors such as smoking, diabetes and hypertension were found to be more prevalent in affected group (25.9%, 33.7% and 45.8) than incontrols (15.7%, 19.6% and 24.7%). Low level of TC (Total cholesterol), LDL (low density lipoprotein) and TG (triglyceride) noted in cad affected group might be due to use of statin in over 80% of affected subjects.

**Table 3.** Baseline characteristics of the study population.

| **Clinical Characteristics** | **Unaffected (N=335)** | **CAD Affected (N=297)** | **P-value** |
| --- | --- | --- | --- |
| Age | 44.57±0.514 | 43.84±0.446 | 0.287 |
| Gender (Male) | 263 (78.5%) | 72 (21.5%) | 0.269 |
| Gender (Female) | 240 (80.8%) | 57 (19.2%) |  |
| Smoking | 52 (15.7%) | 77 (25.9%) | 0.001 |
| Alcohol | 61 (18.5%) | 68 (22.9%) | 0.103 |
| Hypertension | 82 (24.7%) | 136 (45.8%) | 2.11*10<sup>-008</sup> |
| Diabetes | 65 (19.6%) | 100 (33.7%) | 4.34*10<sup>-005</sup> |
| BMI | 25.87±0.288 | 25.59±0.94 | 0.489 |
| Waist Circumference (cm) | 89.86±0.797 | 88.26±0.526 | 0.094 |
| Waist / Hip Ratio | 0.943±0.004 | 0.931±0.007 | 0.011 |
| LDL | 107.04±1.91 | 91.36±2.61 | 1.67*10<sup>-006</sup> |
| HDL | 40.85±0.56 | 37.12±0.56 | 3.46*10<sup>-006</sup> |
| Total Chol. (mg/dl) | 178.54±2.13 | 153.69±2.74 | 2.45*10<sup>-012</sup> |
| Trigly | 158.73±5.43 | 152.76±5.24 | 0.429 |
| LDL/HDL | 2.71±0.05 | 2.17±0.09 | 8.43*10<sup>-007</sup> |
| TC/HDL | 4.51±0.06 | 4.01±0.11 | 0.0002 |
| PhosphoGSK3b | 451.61±8.58 | 571.36±14.25 | 2.33*10<sup>-012</sup> |
| Statin | 13 (4.3%) | 242 (81.8%) | 1.07*10<sup>-094</sup> |
| BetaBlocker | 21 (6.9%) | 211 (71.3%) | 3.43*10<sup>-065</sup> |
| Fibrate | 4 (1.3%) | 2 (0.7%) | 0.356 |
| Calcium channel Blocker | 20 (6.6%) | 44 (14.9%) | 0.001 |
| ACE inhibitor | 6 (2.0%) | 89 (30.1%) | 7.062*10<sup>-024</sup> |
| Antiplatelet | 10 (3.3%) | 252 (85.1%) | 1.04*10<sup>-106</sup> |
| Hypoglycemic agents | 32 (10.5%) | 26 (8.8%) | 0.280 |
| Nitrate | 1 (0.3%) | 73 (24.7%) | 1.00*10<sup>-023</sup> |

Phospho-GSK3B levels in CAD affected population was 571.36±14.25. The control population had phospho-GSK3B levels 451.61±8.58 (p< 0.0001). This difference remained significant even after adjusting for the classical risk factors (hypertension, diabetes, smoking, alcohol) (P<0.0001).

Subset analysis was also performed to check the association of PhopshoGsk3b with individual Risk factor (table 4a):

**Table 4a.** Subset analysis of Classical risk factors

|  |  | **Unaffected** | **CAD Affected** | **P-value** |
| --- | --- | --- | --- | --- |
| **Only Hypertension** | PhosphoGSK3b (pg/ml) | 455.30±16.71 | 506.47±18.90 | 0.44 |
| **Only Diabetes** | PhosphoGSK3b (pg/ml) | 452.76±21.68 | 505±23.07 | 0.100 |
| **Only Smoking** | PhosphoGSK3b (pg/ml) | 455.19±21.03 | 615.97±27.79 | $9.63*10^{-006}$ |
| **Only alcohol** | PhosphoGSK3b (pg/ml) | 523.03±19.79 | 713.38±27.59 | $1.37*10^{-007}$ |

In the subset analysis of hypertension we identified, out of the 632 patients 218 subjects had hypertension of these 136 subjects were from CAD affected group and the phosphoGSK3B level of 506.47±18.91 and remaining 82 control subjects with Hypertension had phosphoGSK3B level of 455.30±16.72 (P<0.05).

In subset analysis of smoking, 129 subjects had smoking in overall subjects of these 77 were from CAD affected and had phosphoGSK3B level of 615.97± 27.79 and remaining 52 control subjects had phspho-GSK3B level of 455.19± 21.04 (P<0.0001).

Among the patients, 68 were alcoholic and had phospho-GSK3B level of 713.38±27.59, whereas in control group 61 were alcoholic and had phspho-GSK3B level of 523.03±19.79 (P<0.0001).

In contrast there were no significant difference were found for phspho-GSK3B with diabetes

### Association of Phspho-GSK3B with CAD

As shown in table 4b, logistic regression analysis using enters method was performed whichrevealed significant association of Phospho-GSK3B with CAD (OR: 4.078, 95% CI: 2.63-6.31, p=2.73*10^−10^). For this model 1, AUC: 0.620 (95% CI: 0.57-0.66) (p= 2.049*10^−7)^ was obtained using ROC analysis. In model 2, The OR increased to 5.65 following adjustment of Traditional risk factors (hypertension, diabetes, smoking and alcohol habit) and the AUC was improved to 0.737 (95%CL: 0.69-0.77, p= 1.12*10^−24^). In model 3, Waist Circumference was added to model2, which revealed highest OR for phspho-GSK3b (OR: 6.27, 95%CI: 3.76-10.44, P= 1.78*10^−12^) and the corresponding AUC was improved to 0.752 (95%CL: 0.71-0.79, P=1.87*10^−27^).

**Table 4b.**
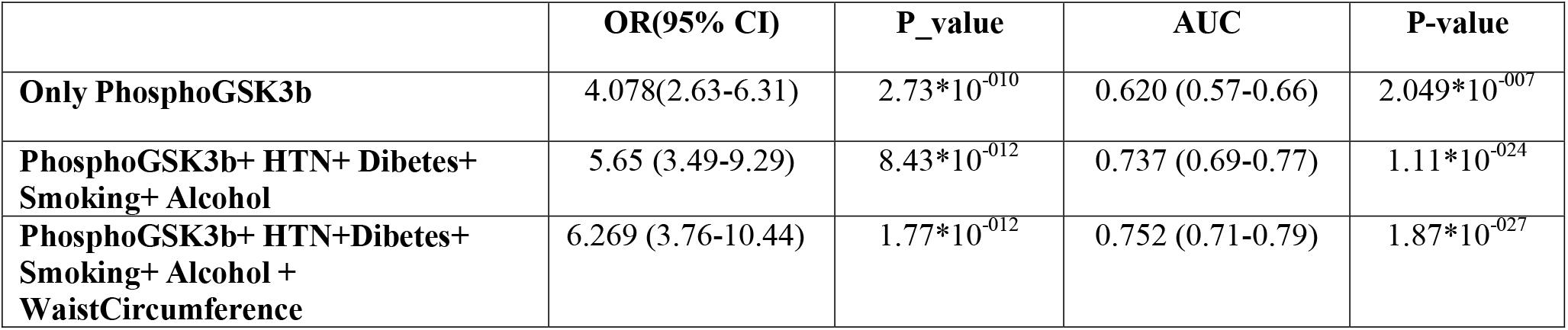
Risk assessment of PhosphoGSK3b

## Discussion

About 43% of phosphorylation sites are there in human genome of which 23% is covered by serine phosphorylation (shown in supplementary table). The same pattern was found in CAD which clearly showed that small change in phosphorylation status may lead to the huge impact on disease. Serine, threonine and tyrosine are most commonly phosphorylation PTMs studied in heart disease. Post translational modifications such as serine and threonine phosphorylation were investigated in cardiac mitochondrial for chronic stimulation of β1-adrenergic receptors/transmembrane AC/cAMP/PKA signaling pathway in heart [25]. Specific group of proteins called protein kinases (PKs) and protein phosphatases (PPs) are responsible for inducing and blocking phosphorylation in downstream proteins which may results in changing its processes and functions. Rapamycin is the target for mTOR, which is a serine/threonine kinase and known to be an important actuator for chronic inflammation. In CAD patients, mTOR phosphorylation induce NF-kB activation leading to p65 nuclear translocation and IL6/TNF-a expression [26]. Another example of protein kinase G (PKG), a serine/threonine kinase which is involved in regulation of intracellular switches that turn on or off the controls of cellular processes like calcium mobilization and phosphorylation of Hsp 20 for instigating vascular dilation. Vasodilator-stimulated phosphoprotein (VASP) is the substrate molecule for PKG and is involved in the regulation of cell migration and motility [27]. GSK3b is one of the important kinase proteins studied in heart disease. GSK3B is involved in several signaling pathways and thus may induce pro and anti-apoptotic roles. Phosphorylation at Ser 9 residue of GSK3b makes it inactivated however, phosphorylation at TYR216 increases its activity. There are conflict reports regarding the pro or anti-atherogenic role of GSK3b. A few studies have shown the pro apoptotic role of active GSK3 by inhibiting prosurvival transcription factors such as CREB and heat shock factor-1. On the other hand, several kinases have been shown to participate in modification of GSK3b, including PKC, AKT and growth factors that work through protein kinase B cascadefor its reducing activity [28-30]. However, literature have also shown that these kinases get regulated by estrogen [31]. Phosphorylated Gsk3b (inactivated Gsk3b) plays an important role in P53 and TNF induced apoptosis [32, 33] through b catenin pathway thatresults in increasedexpression of several inflammatory markers including Ccl24, Cxcl2, Cxcl10, Cxcl11, IL15, IL18, and Ikkβ [34]. Inhibition of GSK3B also increases the transcriptional activity of heat shock protein which further stimulate endothelial cells, macrophages, and SMCs which are most important process in atherosclerosis [35, 36]. In the current study we observed the links between GSK3B, ESR1, HSP1, MYC, JUN, HSF1, AKT1, PRKACA and PRKCB1. Taking this into consideration we hypothesize that due to external stimulus like infection and inflammatory conditions, ESR1 may activate PKA and AKT pathways, in turn inhibiting GSK3B. This Inhibition of GSK3B may increase the transcriptional activity of heat shock protein and promote vascular smooth muscle cell growth through the downstream regulation of c-JUN and MYC (known as oncoproteins) [37, 38] which in turn help in progression of atherosclerotic plaque.

In conclusion, an integrative translation informatics approach has been used in this study for identification of the important PTMs in Disease and its hub regulators. The results of the present study suggest that Serine phosphorylation is highly associated with CAD. Further, PhopshoGSK3b has been identified as best predictive marker for Asian Indian population and contributes maximally to CAD when adjusted with CRFs such as Hypertension, Diabetes, Obesity and Alcohol. However, additional prospective randomized trials are needed and it is necessary to perform biomarker assays in larger population to validate the observations of the current study.

## Abbreviations

CAD: Coronary artery disease
GSK3b: Glycogen synthase kinase 3 beta
PTM: Post translation modification

## ACKNOWLEDGEMENTS

The current study was supported by the Bharati Foundation, India (grant no. 005/2012-2013). Sponsors had no role in the design, conduct, sample collection, analysis and interpretation of the data or in the preparation, review or approval of the manuscript. The authors would like to thank all investigators, staff and administrative teams and participants of IARS at Narayana Institute of Cardiac Sciences, Bangalore (India) and the Asian Heart Institute, Mumbai (India) for their contributions. The authors would also like to thank the patients and their family members for participating in the study.

## Notes

### Competing Interest Statement

The authors have declared no competing interest.

## References

1. Reddy, K.S., et al., Responding to the threat of chronic diseases in India. The Lancet, 2005. 366(9498): p. 1744–1749.

2. Khot, U.N., et al., Prevalence of conventional risk factors in patients with coronary heart disease. Jama, 2003. 290(7): p. 898–904.

3. Falk, E., Pathogenesis of atherosclerosis. Journal of the American College of Cardiology, 2006. 47(8): p. C7–C12.

4. Zhi, Y. and R.M. Sandri-Goldin, Analysis of the phosphorylation sites of herpes simplex virus type 1 regulatory protein ICP27. Journal of virology, 1999. 73(4): p. 3246–3257.

5. Brunet, A., et al., Stress-dependent regulation of FOXO transcription factors by the SIRT1 deacetylase. science, 2004. 303(5666): p. 2011–2015.

6. Yuan, Z.-l., et al., Stat3 dimerization regulated by reversible acetylation of a single lysine residue. Science, 2005. 307(5707): p. 269–273.

7. Vadseth, C., et al., Pro-thrombotic state induced by post-translational modification of fibrinogen by reactive nitrogen species. Journal of Biological Chemistry, 2004. 279(10): p. 8820–8826.

8. Mair, J., Clinical significance of pro-B-type natriuretic peptide glycosylation and processing. 2009, Clinical Chemistry.

9. Kanjilal, S., et al., Prevalence and component analysis of metabolic syndrome: an Indian atherosclerosis research study perspective. Vascular health and risk management, 2008. 4(1): p. 189.

10. Cheng, D., et al., PolySearch: a web-based text mining system for extracting relationships between human diseases, genes, mutations, drugs and metabolites. Nucleic acids research, 2008. 36(suppl 2): p. W399–W405.

11. Plake, C., et al., AliBaba: PubMed as a graph. Bioinformatics, 2006. 22(19): p. 2444–2445.

12. Rebholz-Schuhmann, D., et al., EBIMed—text crunching to gather facts for proteins from Medline. Bioinformatics, 2007. 23(2): p. e237–e244.

13. Health, U.N.I.o., http://ClinicalTrials.gov. 2012.

14. Wishart, D.S., et al., DrugBank: a comprehensive resource for in silico drug discovery and exploration. Nucleic acids research, 2006. 34(suppl 1): p. D668–D672.

15. Liu, H., et al., CADgene: a comprehensive database for coronary artery disease genes. Nucleic acids research, 2011. 39(suppl 1): p. D991–D996.

16. Lee, T.-Y., et al., dbPTM: an information repository of protein post-translational modification. Nucleic acids research, 2006. 34(suppl 1): p. D622–D627.

17. Apweiler, R., et al., UniProt: the universal protein knowledgebase. Nucleic acids research, 2004. 32(suppl 1): p. D115–D119.

18. Minguez, P., et al., Deciphering a global network of functionally associated post_translational modifications. Molecular systems biology, 2012. 8(1): p. 599.

19. Minguez, P., et al., PTMcode v2: a resource for functional associations of post-translational modifications within and between proteins. Nucleic acids research, 2014: p. gku1081.

20. Shannon, P., et al., Cytoscape: a software environment for integrated models of biomolecular interaction networks. Genome research, 2003. 13(11): p. 2498–2504.

21. Marquez, J., et al., Post-translational modifications of cardiac mitochondrial proteins in cardiovascular disease: not lost in translation. Korean circulation journal, 2016. 46(1): p. 1–12.

22. Yang, C.-Y., et al., PhosphoPOINT: a comprehensive human kinase interactome and phospho-protein database. Bioinformatics, 2008. 24(16): p. i14–i20.

23. DeLong, E.R., D.M. DeLong, and D.L. Clarke-Pearson, Comparing the areas under two or more correlated receiver operating characteristic curves: a nonparametric approach. Biometrics, 1988: p. 837–845.

24. Pavlidis, P. and W.S. Noble, Matrix2png: a utility for visualizing matrix data. Bioinformatics, 2003. 19(2): p. 295–296.

25. Rosca, M., P. Minkler, and C.L. Hoppel, Cardiac mitochondria in heart failure. Biochimica et Biophysica Acta-Bioenergetics, 2011. 1807(11): p. 1373–1382.

26. Gao, S., et al., The activation of mTOR is required for monocyte pro-inflammatory response in patients with coronary artery disease. Clinical Science, 2015. 128(8): p. 517–526.

27. Holt, A.W., et al., Soluble guanylyl cyclase-activated cyclic GMP-dependent protein kinase inhibits arterial smooth muscle cell migration independent of VASP-serine 239 phosphorylation. Cellular Signalling, 2016. 28(9): p. 1364–1379.

28. Matsuda, T., et al., Distinct roles of GSK-3α and GSK-3β phosphorylation in the heart under pressure overload. Proceedings of the National Academy of Sciences, 2008. 105(52): p. 20900–20905.

29. Cross, D.A., et al., Inhibition of glycogen synthase kinase-3 by insulin mediated by protein kinase B. Nature, 1995. 378(6559): p. 785.

30. Eldar-Finkelman, H., et al., Inactivation of glycogen synthase kinase-3 by epidermal growth factor is mediated by mitogen-activated protein kinase/p90 ribosomal protein S6 kinase signaling pathway in NIH/3T3 cells. Journal of Biological Chemistry, 1995. 270(3): p. 987–990.

31. Kazi, A.A., K.H. Molitoris, and R.D. Koos, Estrogen rapidly activates the PI3K/AKT pathway and hypoxia-inducible factor 1 and induces vascular endothelial growth factor A expression in luminal epithelial cells of the rat uterus. Biology of reproduction, 2009. 81(2): p. 378–387.

32. Deng, J., et al., β-catenin interacts with and inhibits NF-κB in human colon and breast cancer. Cancer cell, 2002. 2(4): p. 323–334.

33. Watcharasit, P., et al., Direct, activating interaction between glycogen synthase kinase-3β and p53 after DNA damage. Proceedings of the National Academy of Sciences, 2002. 99(12): p. 7951–7955.

34. Anson, M., et al., Oncogenic β-catenin triggers an inflammatory response that determines the aggressiveness of hepatocellular carcinoma in mice. The Journal of clinical investigation, 2012. 122(2): p. 586–599.

35. Asea, A., et al., Novel signal transduction pathway utilized by extracellular HSP70 role of Toll-like receptor (TLR) 2 and TLR4. Journal of Biological Chemistry, 2002. 277(17): p. 15028–15034.

36. Xavier, I.J., et al., Glycogen synthase kinase 3β negatively regulates both DNA-binding and transcriptional activities of heat shock factor 1. Journal of Biological Chemistry, 2000. 275(37): p. 29147–29152.

37. Yada, M., et al., Phosphorylation_dependent degradation of c_Myc is mediated by the F_box protein Fbw7. The EMBO journal, 2004. 23(10): p. 2116–2125.

38. Patel, S. and J. Woodgett, Glycogen synthase kinase-3 and cancer: good cop, bad cop? Cancer cell, 2008. 14(5): p. 351–353.

