## supplementary table 2 for "A systems biology approach to elucidate the post-translational regulome of coronary artery disease"

| No. | PTMs | Whole genome |
| --- | --- | --- |
| 1. | Serine Phosphorylation | 49989 |
| 2. | Threonine Phosphorylation | 16342 |
| 3. | Tyrosine Phosphorylation | 12258 |
| 4. | Acetylation | 7515 |
| 5. | N-linked Glycosylation | 2951 |
| 6. | O-linked Glycosylation | 2169 |
| 7. | ADP-ribosylation | 13 |
| 8. | Alkylation | 7 |
| 9. | Amidation | 46 |
| 10. | Carbamidation | 25 |
| 11. | Carboxylation | 45 |
| 12. | Citrullination | 27 |
| 13. | C-linked Glycosylation | 51 |
| 14. | Deamidation | 20 |
| 15. | Disulfide bond | 1135 |
| 16. | Hydroxylation | 274 |
| 17. | Methylation | 539 |
| 18. | Myristoylation | 81 |
| 19. | Neddylation | 29 |
| 20. | Nitration | 38 |
| 21. | Oxidation | 21 |
| 22. | Palmitoylation | 177 |
| 23. | Prenylation | 51 |
| 24. | Proteolytic Cleavage | 903 |
| 25. | Sumoylation | 604 |
| 26. | Ubiquitylation | 21049 |
| 27. | Others | 1688 |
| 28. | Total | 118047 |

Supplementary table 2: Distribution of PTM in whole genome
